# Chromatin interaction maps identify Wnt responsive *cis*-regulatory elements coordinating *Paupar-Pax6* expression in neuronal cells

**DOI:** 10.1101/2021.05.18.442939

**Authors:** Ioanna Pavlaki, Michael Shapiro, Giuseppina Pisignano, Jelena Telenius, Silvia Muñoz Descalzo, Robert J. Williams, Jim R. Hughes, Keith W. Vance

## Abstract

Central nervous system-expressed long non-coding RNAs (lncRNAs) are often located in the genome close to protein coding genes involved in transcriptional control. Such lncRNA-protein coding gene pairs are frequently temporally and spatially co-expressed in the nervous system and are predicted to act together to regulate neuronal development and function. Although some of these lncRNAs also bind and modulate the activity of the encoded transcription factors, the regulatory mechanisms controlling co-expression of neighbouring lncRNA-protein coding genes remain unclear. Here, we used high resolution NG Capture-C to map the *cis*-regulatory interaction landscape of the key neuro-developmental *Paupar-Pax6* lncRNA-mRNA locus. The results defined chromatin architecture changes associated with high *Paupar-Pax6* expression in neurons and identified both promoter selective as well as shared *cis*-regulatory interactions with the *Paupar* and *Pax6* promoters involved in regulating *Paupar-Pax6* co-expression in neuronal cells. The TCF7L2 transcription factor, a major regulator of chromatin architecture and effector of the Wnt signalling pathway, binds to a subset of these candidate *cis*-regulatory elements to coordinate *Paupar* and *Pax6* co-expression. We identify a functional TCF7L2 bound *cis*-regulatory element within the *Paupar* gene, suggesting that the *Paupar* DNA locus itself regulates *Pax6* expression in addition to its previously described transcript-dependent modes of action. Our work provides important insights into the chromatin interactions, signalling pathways and transcription factors controlling co-expression of adjacent lncRNAs and protein coding genes in the brain.

## Introduction

A typical gene promoter is regulated by multiple different types of *cis*-regulatory elements (CREs) such as transcriptional enhancers and silencers. These are DNA sequences containing clusters of transcription factor binding sites that act together to generate the correct temporal and spatial expression of their target genes (Jeziorska, Jordan et al. 2009). Chromatin conformation capture (3C) based technologies have shown that short- and long-range dynamic chromatin looping interactions bring CREs and their target promoters into close physical proximity in the nucleus to facilitate gene regulation. More recently, high throughput 3C variants such as NG Capture-C have been used to map large numbers of CREs to their cognate genes at high resolution and investigate the complexity of CRE-promoter communication at unprecedented detail (Davies, Telenius et al. 2016, Davies, Oudelaar et al. 2017).

Precise temporal and spatial control of expression of the *Pax6* transcription factor gene is required for the normal development and function of the nervous system. *Pax6* haploinsufficiency in mice results in abnormal eye and nasal development and causes a range of brain defects; whilst mutations affecting *PAX6* expression and function in humans cause anirida, an autosomal dominant inherited disorder characterized by a complete or partial absence of the iris (Jordan, Hanson et al. 1992, Manuel, Mi et al. 2015, Lima Cunha, Arno et al. 2019). *Pax6* is transcribed from 2 major upstream promoters (P0, P1) and multiple CREs have been shown to control *Pax6* expression in distinct domains in the central nervous system and eye (Williams, Altmann et al. 1998, Xu, Zhang et al. 1999, Morgan 2004). These include the neuroretina, ectodermal and retinal progenitor enhancers just upstream of the P0 promoter (Kammandel, Chowdhury et al. 1999, Plaza, Saule et al. 1999); a photoreceptor enhancer situated between the P0 and P1 promoters (Xu, Zhang et al. 1999); the retina regulatory region located between exons 4 and 5 (Kammandel, Chowdhury et al. 1999, Xu, Zhang et al. 1999); and three conserved sequence elements within intron 7 that activate *Pax6* in the diencephalon, rhombencephalon and at late stages of eye development (Kleinjan, Seawright et al. 2004). These CREs are all located within a 30 kb window surrounding the *Pax6* P0 and P1 promoters and act over short genomic distances. In addition, several candidate long-range enhancers have been identified approximately 150-200 kb downstream of the *Pax6* gene and some of these have also been shown to drive *Pax6* expression in specific domains of the eye and brain (Kleinjan, Seawright et al. 2006, McBride, Buckle et al. 2011). However, these enhancers together are not sufficient to generate the full temporal and spatial pattern of *Pax6* expression in the central nervous system suggesting the presence of additional uncharacterised *Pax6* regulatory elements.

Thousands of lncRNAs are temporally and spatially expressed within the central nervous system and some of these are thought to be important in brain development and function (Derrien, Johnson et al. 2012, Liu, Nowakowski et al. 2016, Hezroni, Ben-Tov Perry et al. 2020). Brain-expressed lncRNAs are preferentially located in the genome close to protein coding genes involved in transcriptional control (Ponjavic, Oliver et al. 2009). This includes bidirectional lncRNAs that are transcribed in the opposite direction to a protein coding gene from a shared promoter as well as intergenic lncRNAs that are either expressed from their own promoter or from a transcriptional enhancer. Such lncRNA-mRNA pairs are frequently co-expressed during neuronal development and in different brain regions and can function in the control of similar biological processes (Chalei, Sansom et al. 2014, Vance, Sansom et al. 2014). The lncRNA *Paupar,* transcribed from a promoter approximately 8.5 kb upstream of the *Pax6* gene, is an important regulator of neurogenesis *in vivo* in mouse and human, and is co-ordinately expressed with *Pax6* during neural differentiation *in vitro* and in the adult mouse brain (Vance, Sansom et al. 2014, Pavlaki, Alammari et al. 2018, Xu, Xi et al. 2021). Moreover, *Paupar* transcript directly binds PAX6 and acts as a transcriptional cofactor to promote the formation of a PAX6-*Paupar*-KAP1 chromatin regulatory complex at important neuronal genes (Vance, Sansom et al. 2014, Pavlaki, Alammari et al. 2018). Even though *Paupar* and *Pax6* can act together to regulate shared biological processes important for neuronal development, the CREs controlling *Paupar-Pax6* co-expression in the nervous system are not known.

Here we used NG Capture-C to generate high resolution chromatin interaction maps with the *Paupar* and *Pax6* promoters in *Paupar-Pax6* high- and low-expressing cells. The results identified shared chromatin interactions with both the *Paupar* and *Pax6* promoters involved in regulating *Paupar-Pax6* co-expression, as well as promoter specific *cis*-regulatory interactions. We show that the Wnt-Bmp4 signalling axis acts through the TCF7L2 transcription factor to co-ordinate *Paupar* and *Pax6* coexpression in neuronal cells, including through a functional TCF7L2 bound *cis*-regulatory element within the *Paupar gene. We* report cell type specific differences in both local and distal chromatin interactions with the *Paupar* and *Pax6* promoters that may be important for CRE-promoter communication and *Paupar-Pax6* activation in neurons. Our work further refines the complex *cis*-regulatory landscape surrounding the *Paupar-Pax6* locus and provides critical insights into the regulatory mechanisms controlling the co-expression of adjacent lncRNAs and protein coding genes in the brain.

## Results

### Identification of cis-regulatory interactions with the Paupar and Pax6 promoters using high resolution NG Capture-C

We previously showed that the lncRNA *Paupar* and its adjacent *Pax6* transcription factor gene are highly expressed in the adult mouse brain, and that *Paupar* and *Pax6* expression are temporally coordinated during *in vitro* neural differentiation of mouse embryonic stem cells (ESCs) (Vance, Sansom et al. 2014). In this study, we used the following neuronal cell types to investigate *Paupar-Pax6* expression control: primary neural stem cells (NSCs) isolated from E14.5 mice, differentiated mouse cortical neurons, N2A mouse neuroblastoma cells, as well as mouse ESCs as a non-neuronal reference. RT-qPCR analysis demonstrated that *Paupar* and *Pax6* P0 and P1 expression is significantly higher in neuronal cell types compared to ESCs, with highest expression in NSCs and differentiated neurons (Fig. 1A, B). Consistent with an earlier report (Kleinjan, Seawright et al. 2001), our results also suggest that the *Pax6* P1 promoter is the major *Pax6* promoter in the neuronal lineage. We found that Pax6 P1 promoter transcription was 40- and 105-fold more active than P0 in NSCs and differentiated neurons respectively, whilst in N2A cells Pax6 P0 expression was undetectable (Fig. 1A, B).

**Figure 1.**
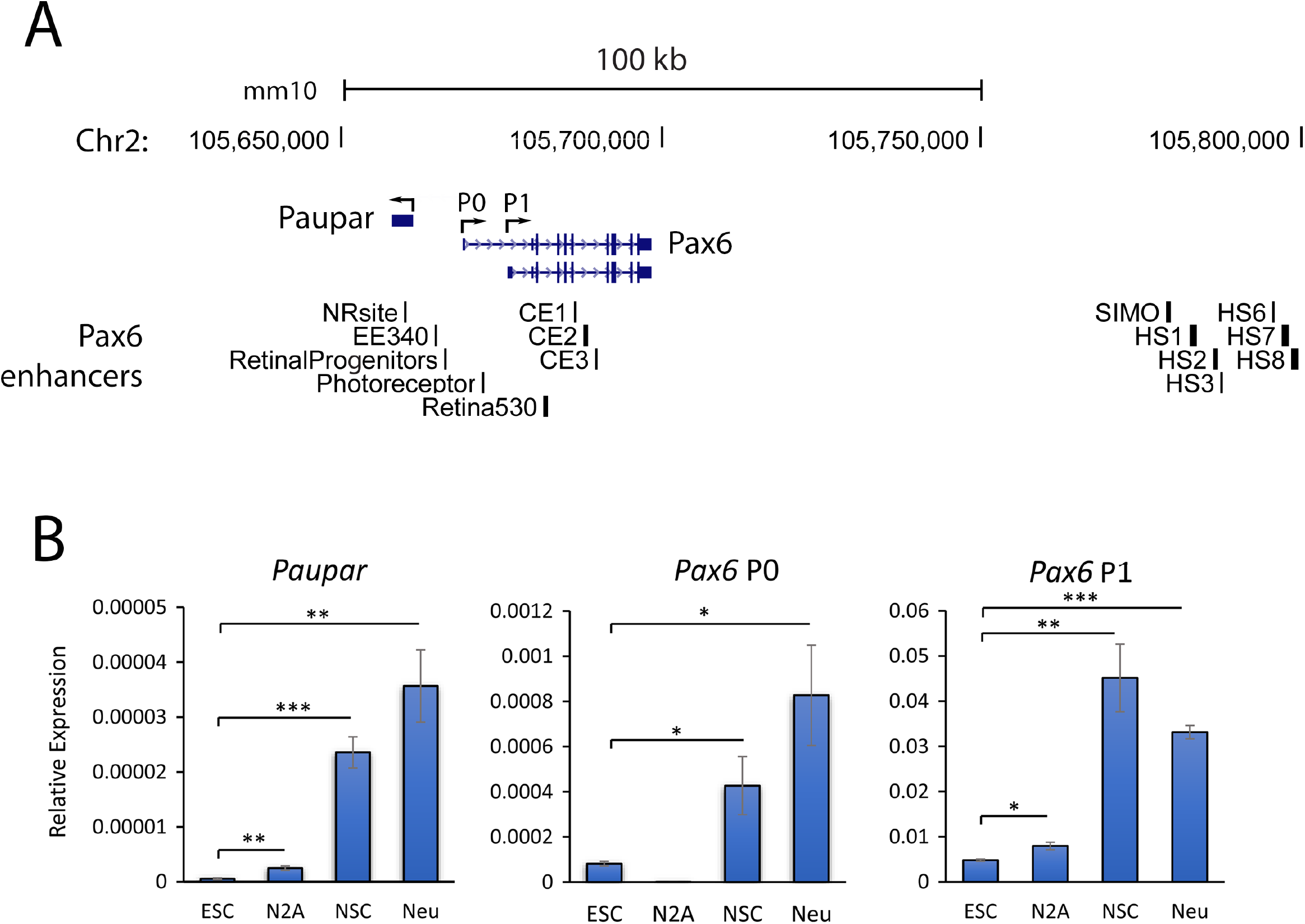
*Paupar* and *Pax6* are co-ordinately expressed to higher levels in neuronal cells compared to ESCs. (A) Genome browser graphic (GRCm38/mm10) showing the location of *Paupar* and *Pax6* genes, the two major *Pax6* promoters (P0 and P1) and known *Pax6* enhancers. NR (neuroretina), EE (ectodermal enhancer), RP (retinal progenitors), PR (photoreceptor), Re (retina), CE1-3 (conserved element 1-3), HS (hypersensitivity site). (B) Transcripts generated from the *Paupar* and *Pax6* P0 and P1 promoters were measured in ESCs, NSCs, differentiated neurons and N2A cells using RT-qPCR. Results are presented relative to the *Tbp* reference gene. Mean values +/− sem shown, n=6, one-tailed t-test, unequal variance. * p<0.05, ** p<0.01, *** p<0.001.

High resolution NG Capture-C was performed to map chromatin interactions with the *Paupar* and *Pax6* P0 and P1 promoters, as well as the Sox2 promoter as a positive control, to identify the *cis*-regulatory DNA sequences important for *Paupar-Pax6* expression control and the chromatin changes associated with activation of the locus in neuronal cells. Multiplexed NG Capture-C libraries were generated and sequenced to an average depth of 61 million paired end reads per library. Benchmarking NG Capture-C data quality using CCanalyser (Davies, Telenius et al. 2016) showed that an average of 37.6% mapped reads contained capture bait sequence across all NG Capture-C libraries, demonstrating good capture enrichment, and confirmed good ligation efficiency as 29.3% of captured fragments were ligated to a reporter (S1 Table). This enabled us to generate high resolution NG Capture-C interaction profiles (Fig. 2A, B, C, S1 Fig) using an average of 33,719 (S2 Table) unique interactions between DpnII restriction fragments and each promoter bait fragment per cell type.

**Figure 2.**
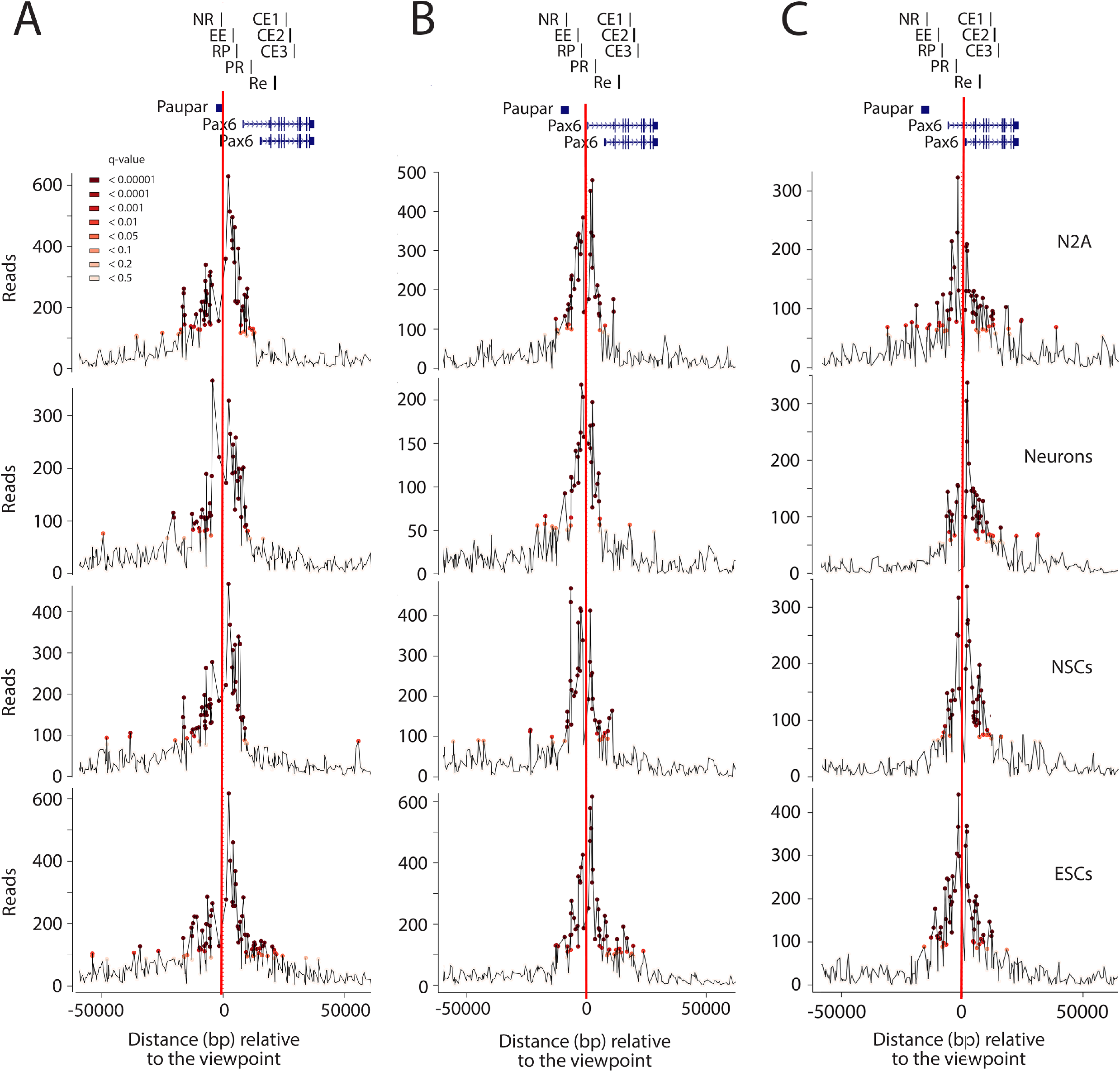
High resolution *cis*-regulatory interactions with the *Paupar* and *Pax6* promoters in *Paupar-Pax6* high- and low-expressing cells. NG Capture-C profiles displaying the interaction count per DpnII restriction enzyme fragment for each NG Capture-C library. The red vertical lines indicate the locations of captured viewpoints. (A) *Paupar* promoter, (B) *Pax6* P0 and (C) *Pax6* P1 promoter viewpoints. Significant fragments determined using R3C-seq (Thongjuea, Stadhouders et al. 2013) are denoted by coloured circles.

NG Capture-C interaction profiles were then analysed using r3C-seq tools (Thongjuea, Stadhouders et al. 2013) to normalise for distance from the capture point and model statistically significant (q < 0.1) CRE-promoter interactions. As expected, we identified significant looping interactions between the *Sox2* super-enhancer overlapping the *Peril* locus (Li, Rivera et al. 2014) and the *Sox2* promoter in ESCs which were not present in neuronal cell types (S1 Fig). This is consistent with previous 3C based maps defining Sox2 enhancer-promoter communication (Li, Rivera et al. 2014) and confirms the ability of our approach to identify functional CREs. To define the set of regulatory interactions mediating *Paupar-Pax6* expression we then determined the number of statistically significant chromatin interactions with the *Paupar* and *Pax6* P0 and P1 promoters present in both biological replicates for each cell type. We discovered that 96% high resolution chromatin interactions are located within a 50 kb window centred around each promoter (Fig 3A), including both upstream and downstream *cis*-acting DNA sequences. The *Paupar-Pax6 cis*-regulatory interaction map showed significant interactions with known *Pax6* regulatory elements as well as many additional short-range regulatory interactions with candidate new CREs involved in *Paupar-Pax6* expression control (Fig 3B and S3 Table for fragment coordinates). Consistent with a role in neuronal gene expression, chromatin interactions with the *Paupar* and *Pax6* promoters show an increased overlap with H3K4me1 ChIP-seq peaks in E12.5 mouse forebrain tissue compared to ESCs using publicly available ENCODE data (He, Hariharan et al. 2020) (Fig 3B). The interaction map also encompasses associations between DNA sequence elements within the *Paupar* genomic locus and the *Pax6* promoters (Fig 3B and S4 Table). Most chromatin interactions (119 out of 168) were found in all cell types tested but we also discovered a subset of neuronal (30/168) and ESC (19/168) specific interactions that may be important for tissue specific regulation of the locus (S5 Table). Our results revealed shared interactions with more than one promoter region that may be important for *Paupar-Pax6* co-expression in the brain (Fig 3C). In addition, we found a subset of specific *cis*-regulatory interactions with individual *Paupar, Pax6 P0* and *P1* promoter viewpoints, suggesting that *Paupar* and *Pax6* expression control may be decoupled (Fig 3C), as well as a small number of trans-interactions with DNA sequences on different chromosomes (Fig 3B and S3 Table). Altogether, these results identify the chromatin interactions and CREs that are likely to be important for precise *Paupar-Pax6* expression control.

**Figure 3.**
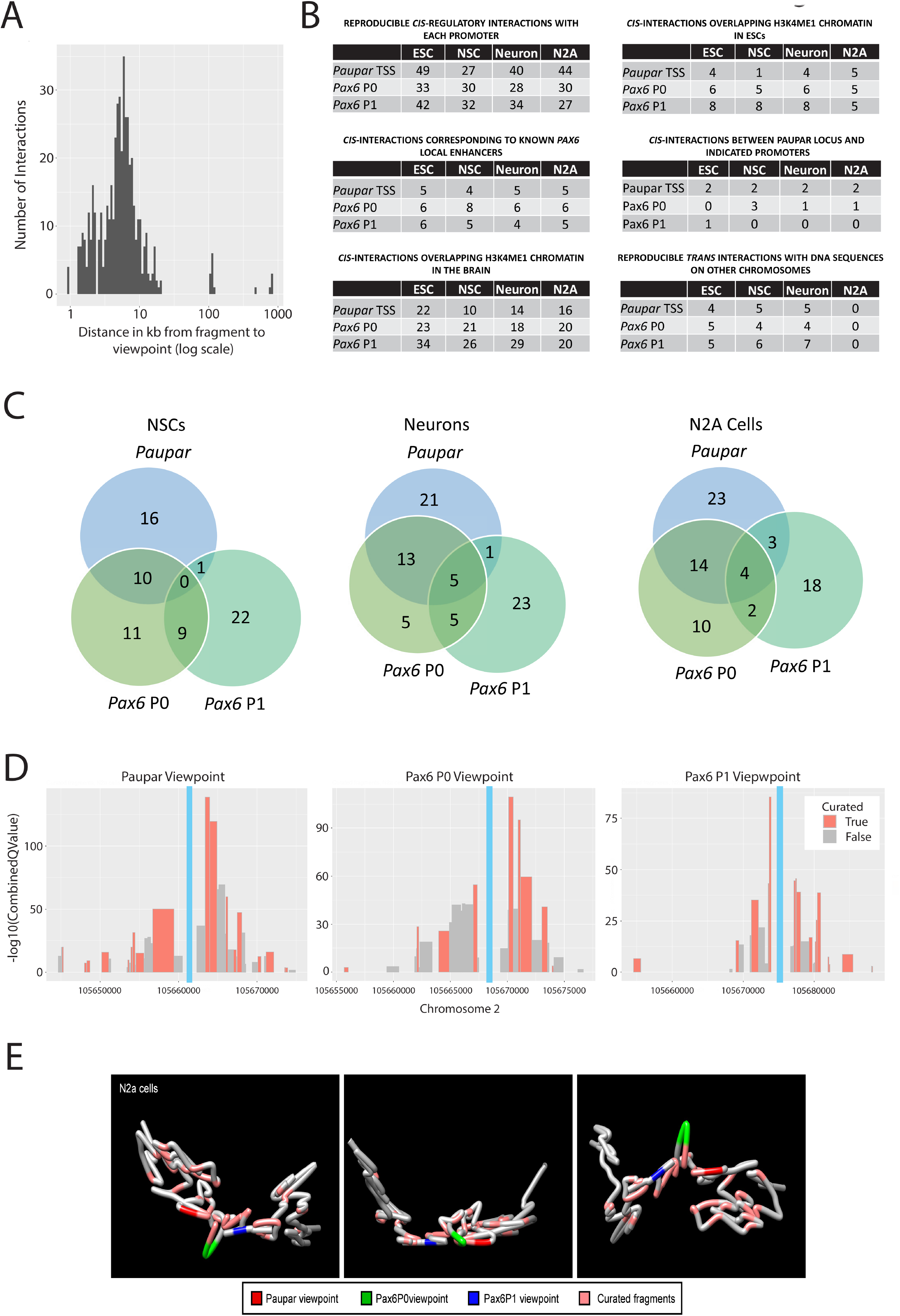
Identification of short-range regulatory interactions with candidate new CREs involved in *Paupar-Pax6* expression control. (A) Histogram depicting distances of *cis*-interacting fragments from their viewpoints. Significant duplicated fragments from all cell types and viewpoints are shown. (B) Tables showing the number of statistically significant reproducible interactions with the indicated promoter viewpoints as well as overlap with known *Pax6* enhancers and enhancer-like chromatin marks using ENCODE data (He, Hariharan et al. 2020). (C) Venn diagram displaying the number of shared and specific interactions with the *Paupar, Pax6* P0 and *Pax6* P1 promoters in NSCs (left), neurons (middle) and N2A cells (right). (D) Curation of interacting fragments for the indicated viewpoints based on combined – log 10 q value from NG Capture-C N2A cell data. q values from replicates were merged by taking the square root of their product. Each bar represents a fragment with significant interaction (q < 0.05). (E) 3D model of *Paupar-Pax6* local chromatin architecture generated from NG Capture-C N2A cell data using the 4Cin software package (Irastorza-Azcarate, Acemel et al. 2018). Viewpoints and curated fragments are marked using the designated colours. The model is shown from three angles.

### Wnt-Bmp4 signalling acts through TCF7L2 to regulate co-expression of the Paupar-Pax6 locus

*We* hypothesized that a subset of discovered genomic fragments would play a causal role in the formation of chromatin interactions needed for expression of the the *Paupar-Pax6* locus and that these would have an elevated discovery rate in our analysis. N2A cells were used to identify and define the function of such sequences as these cells represent a well characterised tractable *in vitro* model of neuronal differentiation and were previously used to determine *Paupar* and *Pax6* gene regulatory functions (Vance, Sansom et al. 2014, Pavlaki, Alammari et al. 2018). To identify sequences with increased proximity to the promoter viewpoints compared to surrounding sequence, we plotted the mean -log10 q-value for each reproducible *cis*-regulatory interaction against chromosome position using N2A cell NG Capture-C data. We next defined the DpnII fragments with the highest local statistical significance and curated a subset of 42 unique fragments (24 for *Paupar,* 10 for *Pax6* P0 and 22 for the *Pax6 P1* viewpoint), visualised as peaks of increased statistical prevalence (Fig 3D). We then applied 4Cin (Irastorza-Azcarate, Acemel et al. 2018) to generate a 3D model of *Paupar-Pax6* local chromatin architecture from N2A cell NG Capture-C data and visualise the relative proximity of the curated fragments to the *Paupar* and *Pax6* promoters. The results showed that most curated fragments are located at curvature points on the chromatin fibre and appear to be orientated towards the *Paupar-Pax6* promoters (Fig 3E). This subset of curated NG Capture-C fragments may represent candidate *Paupar-Pax6* CREs with roles in the regulation of short-range chromatin interactions.

CRE-promoter communication is mediated by protein-protein interactions between transcription factors bound to specific motifs within CREs and proteins assembled at the target promoters. To investigate the transcription factors controlling *Paupar-Pax6* co-expression in the brain we used the Bi Fa motif discovery tool (Jeziorska, Koentges et al. 2012) to search for transcription factor position frequency matrices (PFMs) within the N2A cell curated fragment dataset and identify putative binding sites for factors with known functions in neuronal development. This discovered 13 high scoring PFMs for the TCF7L2/TCF4 transcription factor (Table S6), an important regulator of chromatin structure and major effector of the Wnt/β-catenin signalling pathway. To explore TCF7L2 regulation of *Paupar-Pax6* we first determined whether Wnt signalling regulates expression of the *Paupar-Pax6* locus. RT-qPCR analysis showed that ectopic expression of a constitutively active β-catenin S33Y protein (Morin, Sparks et al. 1997) significantly reduced both *Paupar* and *Pax6* expression by 40% and 33% respectively in N2A cells (Fig 4A). In addition, activation of canonical Wnt signalling using recombinant WNT3a ligand led to a significant 39% reduction in *Tcf7l2* expression and a concomitant 56% reduction in *Paupar* and 47% reduction in *Pax6* levels in N2A cells after 72 hours (Fig 4B). As crosstalk between the Wnt and Bmp signalling pathways is important during neuronal development we then treated N2A cells with BMP4, a *Wnt3a* target gene in neuroblastoma (Szemes, Melegh et al. 2020). This led to a 2.34-fold increase in *Tcf7l2* and a subsequent 2.13- and 1.89-fold up-regulation in *Paupar* and *Pax6* expression after 72 hours (Fig 4C). Altogether, these results suggest that the Wnt-Bmp signalling axis acts through *Tcf7l2* to coordinately regulate both *Paupar* and *Pax6* expression in neural cells.

**Figure 4.**
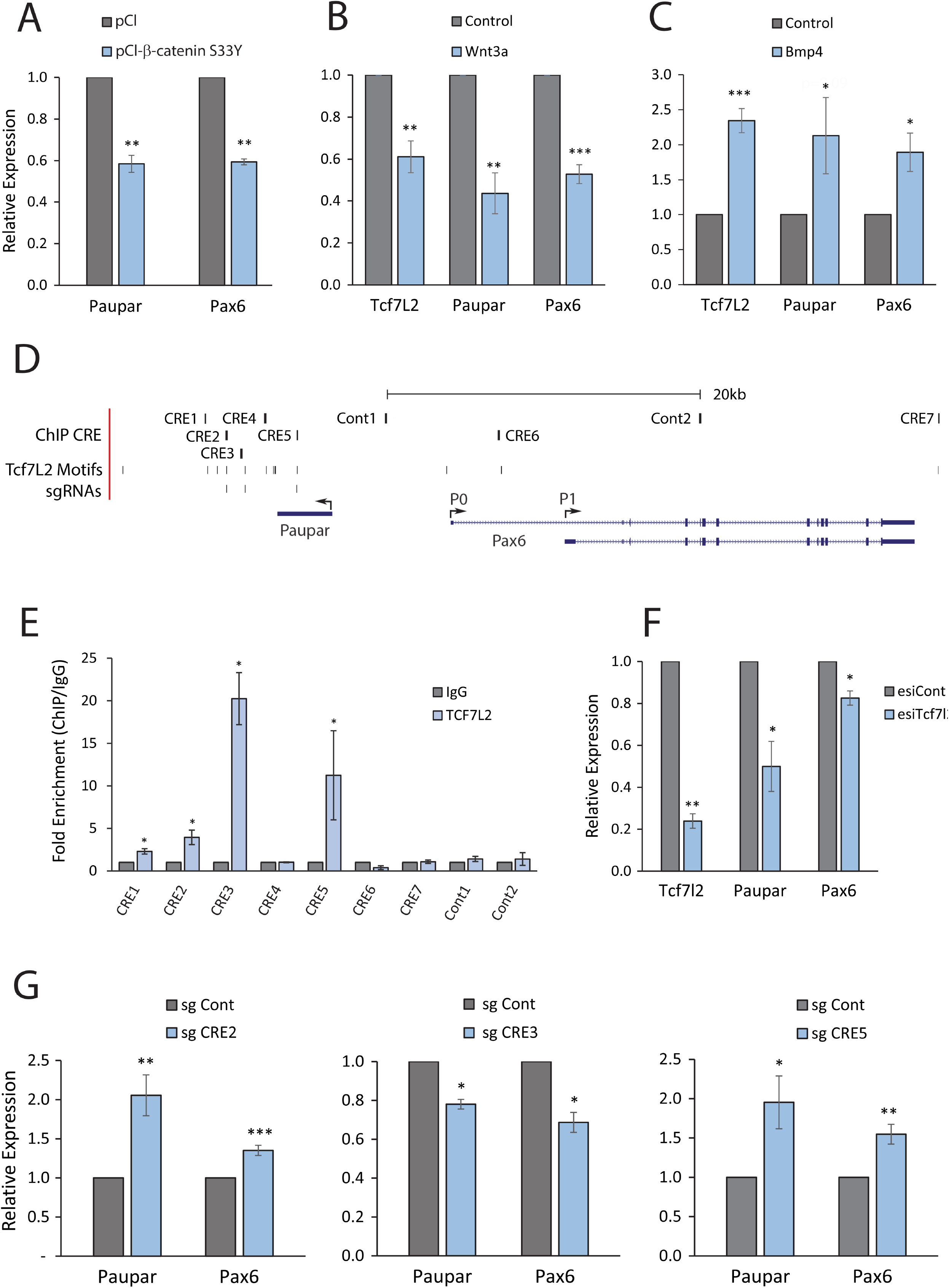
Wnt-Bmp signalling acts through TCF7L2 to coordinate *Paupar-Pax6* expression in neuronal cells. (A) N2A cells were transfected with pCl β-catenin S33Y or pCl empty vector and harvested for expression analysis 2 days later. Results are presented relative to the pCl control. (B, C) N2A cells were treated with either 50 ng/μl Wnt3a (B) or 0.1 ng/μl BMP4 (C) and harvested for RT-qPCR analysis 3 days later. 0.02% BSA in PBS was used as a negative control. (D) Genome browser graphic showing the ChIP amplified candidate CREs as well as the location of the TCF7L2 motifs and sgRNAs used in the CRISPRi experiment (GRCm38/mm10). (E) ChIP assays were performed in N2A cells using either an antibody against TCF7L2 or an isotype specific control. TCF7L2 occupancy at the indicated CREs was analysed by qPCR. Fold enrichment was calculated as 2^−ΔΔCt^ (IP/IgG) and is presented as mean value +/− sem. n≥3. One-tailed student’s t-test * p<0.05. (F) TCF7L2 levels were knocked-down in N2A cells by transfection of an endoribonuclease-prepared pool of TCF7L2 esiRNAs. An esiRNA pool targeting the luciferase gene was used as a negative control. Cells were harvested for expression analysis 3 days after transfection. (G) N2A cells were transfected with a plasmid co-expressing dCas9-KRAB and a sgRNA targeting the TCF7L2 motif within the indicated candidate CREs. A non-targeting sgRNA was used as a control. For all RT-qPCR reactions: *Tcf7l2, Paupar* and *Pax6* expression levels were measured using RT-qPCR and results were normalised to *Tbp.* Results are presented as mean values +/− sem, n≥3. One-tailed student’s t-test * p<0.05, ** p<0.01, *** p<0.001.

### Novel TCF7L2 bound CREs control Paupar-Pax6 co-expression

We next investigated whether *Tcf7l2* directly regulates *Paupar-Pax6.* To do this, ChIP-qPCR was performed to test whether TCF7L2 binds to any of its predicted motifs within the curated fragment dataset. We amplified 7 candidate CREs (CRE1-7) close to TCF7L2 motifs (Fig 4D) and identified strong (> 3-fold enrichment) TCF7L2 chromatin binding at three of them in N2A cells. TCF7L2 binding was 4-fold enriched at CRE2; 20-fold enriched at CRE3; and 11-fold enriched at CRE5, compared to an IgG isotype control (Fig 4E). We next tested whether TCF7L2 regulates *Paupar* and/or *Pax6* expression. TC7FL2 levels were depleted in N2A cells using an endoribonuclease-prepared pool of TCF7L2 siRNAs (esiRNAs) and changes in *Paupar* and *Pax6* expression measured using RT-qPCR (Fig 4F). The results showed that silencing *Tcf7l2* by approximately 76% resulted in a significant 50% reduction in *Paupar* and 18% reduction in *Pax6* expression. Taken together, these results suggest that TCF7L2 binds a subset of DNA sequence elements within NG Capture-C identified candidate CREs to directly activate both *Paupar* and *Pax6* expression.

We then performed CRISPR interference (CRISPRi) to study the function of the TCF7L2 binding sites within their endogenous chromatin context in *Paupar-Pax6* expression control. To do this, a single guide RNA (sgRNA) targeting a specific TCF7L2 motif was used to recruit a catalytically inactive dCas9-KRAB fusion protein to induce local chromatin closing and block regulatory element activity in N2A cells (Fulco, Munschauer et al. 2016). RT-qPCR analysis of *Paupar* and *Pax6* expression showed that targeting dCas9-KRAB to CRE3 induced a significant 22% reduction in *Paupar* and a 31% reduction in *Pax6* expression (Fig 4G, middle panel). CRISPRi against the TCF7L2 motif in CRE2 led to a 2.1-fold up-regulation of *Paupar* and a 1.4-fold increase in *Pax6* expression (Fig 4G, left panel); whilst inhibition of CRE5, located within the *Paupar* DNA locus, resulted in a 2.0-fold increase in *Paupar* and a 1.5-fold increase in *Pax6* expression (Fig 4G, right panel). These results suggest that the CRE3 TCF7L2 motif functions as part of a shared transcriptional enhancer of both *Paupar* and *Pax6,* and that the CRE2 and CRE5 TCF7L2 motif containing sequences co-ordinately repress both *Paupar* and *Pax6.* The *Paupar* DNA locus therefore appears to play a regulatory role in *Pax6* expression control.

### Identification of cell type specific chromatin architecture changes associated with high Paupar and Pax6 expression in neurons

Comparative analyses of NG Capture-C profiles have previously been used to investigate *cis*-regulatory mechanisms controlling cell type specific gene expression (Davies, Telenius et al. 2016). We thus developed a new statistical method to compare the NG Capture-C profiles from *Paupar-Pax6* high-expressing differentiated neurons and low-expressing ESCs and detect changes in chromatin architecture associated with increased *Paupar-Pax6* expression in the brain. In this, normalised NG Capture-C data were first grouped into bins of discrete sizes to increase signal over noise. The results revealed that a 10 kb bin size facilitated the identification of changes in chromatin conformation due to clustering of neighbouring fragments that were not significant at the individual fragment level (S2 Fig). Furthermore, these changes were not discovered using permuted data validating specificity.

Examination of changes in local chromatin architecture surrounding the *Paupar-Pax6* locus (+/− 50kb from each viewpoint) identified an increase in chromatin interactions upstream of the *Pax6* gene with the *Paupar* promoter and an increase in downstream interactions with the *Pax6* P1 promoter in neurons compared to ESCs, as well as a set of reciprocal interactions that increase in ESCs compared to neurons (Fig 5A). Consistently, permutation testing to compare the frequency of upstream versus downstream chromatin interactions from each viewpoint revealed significant asymmetry in chromatin architecture (Fig 5A). These changes in local chromatin organisation may be important for the rewiring of short-range CRE-promoter interactions controlling *Paupar-Pax6* co-expression in neurons, and are consistent with ENCODE ChIP-seq data (He, Hariharan et al. 2020) showing that the chromatin surrounding the *Paupar* and *Pax6* promoters is marked by an increase in open (H3K4me1, H3K4me3, H3K27ac) chromatin marks in E12.5 mouse forebrain tissue compared to ESCs (Fig 5B and S3 Fig).

**Figure 5.**
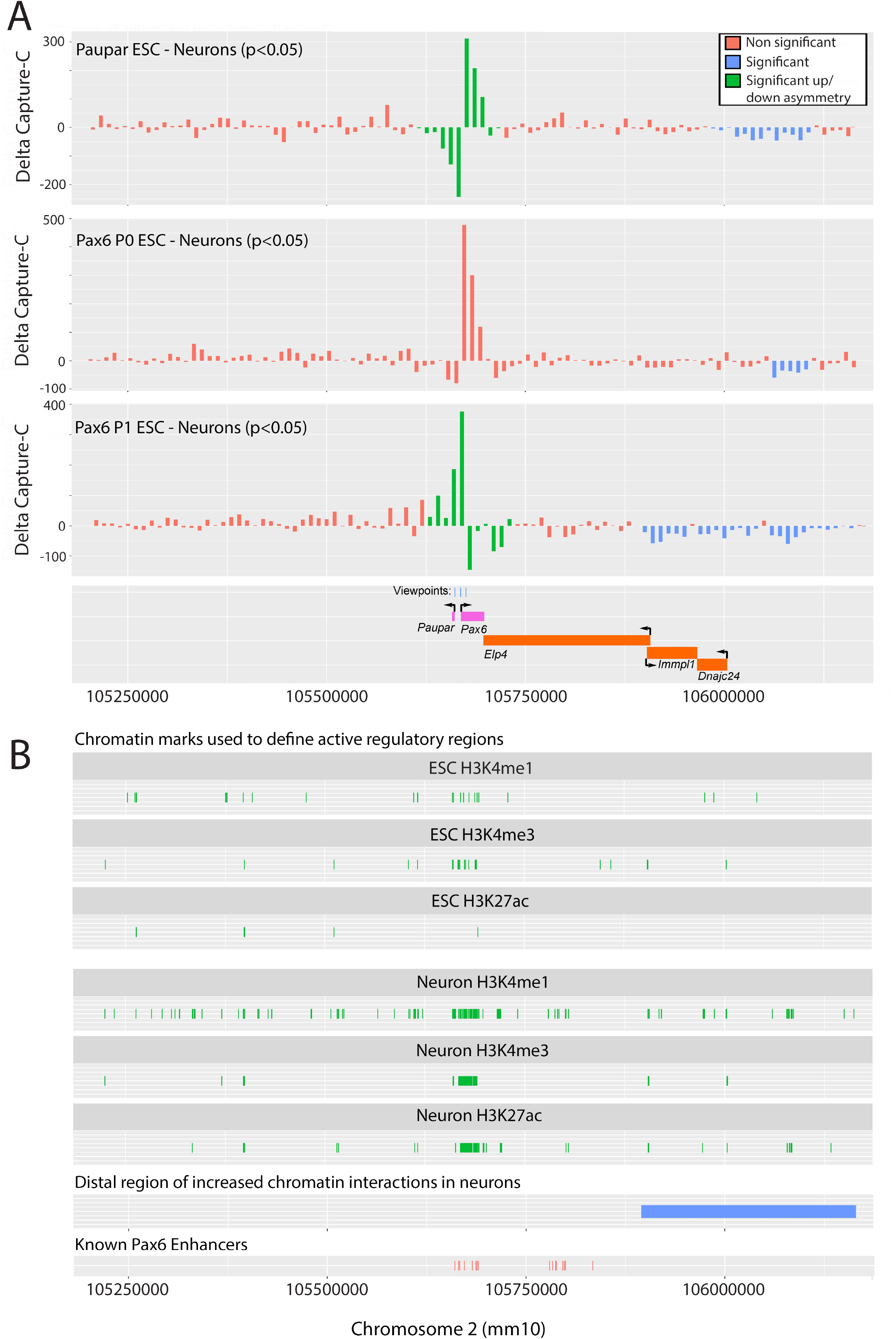
Local and distal chromatin changes associated with increased *Paupar-Pax6* expression in neuronal cells. (A) A comparative analysis of NG Capture-C data to map changes in chromatin conformation between cell types. Differences in mean normalized capture counts for interactions with the *Paupar* (top), *Pax6* P0 (middle) and *Pax6* P1 (bottom) viewpoints between ESC and neurons are plotted on the y-axis. X-axis shows position on chromosome 2 (GRCm38/mm10). Data are binned to 10 kb across approximately ±500 kb genomic sequence around each viewpoint. Permutation testing was performed to determine statistical significance as described in Materials and Methods. (B) ENCODE Project ChIP-seq data mapping the location of H3K4me1, H3K4me3 and H3K27ac peaks in ESCs and mouse forebrain tissue across approximately 1MB genomic sequence surrounding the *Paupar-Pax6* locus (He, Hariharan et al. 2020). Individual peaks of less than 1 kb are shown at 1 kb long for visibility reasons.

As transcriptional regulatory elements can function over large genomic distances, we next analysed 1 MB genomic sequence surrounding the *Paupar-Pax6* locus using binned NG Capture-C data to map meso-scale changes in chromatin architecture between cell types. This detected a large up to 250 kb chromosomal region located approximately 350 kb downstream of the *Pax6* gene that contains an increased frequency of statistically significant interactions with the *Paupar* and *Pax6* promoters in differentiated neurons compared to ESCs (Fig 5A). This region corresponds to an equivalent region in the human genome containing multiple predicted long-range CREs that loop onto the human Pax6 promoter in Promoter Capture Hi-C experiments (Freire-Pritchett, Schoenfelder et al. 2017). Furthermore, this region contains an increased number of H3K4me1, H3K4me3 and H3K27ac ChIP-seq peaks (He, Hariharan et al. 2020) in mouse forebrain tissue compared to ESCs (Fig 5B and S3 Fig). As these histone modifications are known to mark active regulatory regions, we predict that this distal domain may contain additional clusters of uncharacterised long-range CREs involved in *Paupar-Pax6* expression control in neuronal cells. Taken together, these data define the local and distal chromatin changes associated with *Paupar-Pax6* expression in neurons.

## Discussion

LncRNAs involved in brain development are frequently co-expressed with their adjacent protein coding genes. These lncRNA-mRNA pairs often function in the same biological processes and some CNS expressed lncRNAs modulate both the expression and transcriptional activity of their neighbouring protein coding genes. A greater understanding of the complex regulatory relationship controlling the expression and function of lncRNA-mRNA pairs in the brain is needed to further define their role in neuronal development and function.

In this study we used high resolution NG Capture-C to comprehensively define chromatin interactions important for *Paupar-Pax6* expression control. Our work revealed an intricate network of short-range *cis*-regulatory interactions with the *Paupar* and *Pax6* P0 and P1 promoters, including interactions with the previously characterised *Pax6* ectodermal, neuroretina, retinal progenitor and photoreceptor enhancers (Kammandel, Chowdhury et al. 1999, Plaza, Saule et al. 1999, Xu, Zhang et al. 1999), as well as many candidate new CREs. The results classified a subset of shared short-range chromatin interactions with both the *Paupar* and *Pax6* promoters that are likely to be involved in regulating *Paupar-Pax6* co-expression in the brain. We detected significant asymmetry in local chromatin architecture surrounding the *Paupar* and *Pax6* P1 promoters and discovered a subset of short-range chromatin interactions with these promoters that increase in differentiated neurons compared to ESCs. We hypothesize that these chromatin changes are important for CRE-promoter communication and cell type specific expression of the locus.

Our results curated a subset of NG Capture-C fragments based on increased local statistical significance that may be central mediators of short-range CRE-promoter interactions. We discovered that TCF7L2 binds a subgroup of these fragments and acts to integrate signals from the *Wnt* and *Bmp* signalling pathways to control *Paupar* and *Pax6* co-expression. Interplay between the *Wnt* and *Bmp* pathways is critical for proper development of the nervous system and in the *Paupar-Pax6* expressing subventricular zone (SVZ) NSC niche has been shown to regulate postnatal NSC selfrenewal and SVZ neurogenesis (Pavlaki, Alammari et al. 2018, Al-Dalahmah, Campos Soares et al. 2020, Al-Dalahmah, Nicholson et al. 2020). Furthermore, the *Wnt-Bmp* signalling axis also promotes growth suppression and differentiation in neuroblastoma (Szemes, Melegh et al. 2020). *TCF7L2* is a key *Wnt* effector in the brain and is required for the production of *Pax6* expressing neural progenitor cells in the neocortex (Chodelkova, Masek et al. 2018). We expect that TCF7L2 acts as an important regulator of *Paupar-Pax6* chromatin organisation and CRE-promoter communication. Accordingly, TCF7L2 silencing leads to genome-wide changes in chromatin architecture and enhancer-promoter interactions in pancreatic and colon cancer cells, whilst TCF-bound *Wnt* responsive enhancers regulate chromatin looping and activation of the *Myc* gene in colorectal cancer (Yochum, Sherrick et al. 2010, Gerrard, Wang et al. 2019, Brown, Dotson et al. 2021). Furthermore, ENCODE data shows that TCF7L2 associates with more than 40% of active enhancers in the genome in different cell lines suggesting that TCF7L2 is a critical regulator of cell-type specific CRE function (ENCODE 2019). Our finding that *Wnt3a-Bmp4* acts through TCF7L2 to co-ordinate *Paupar-Pax6* co-expression in neuronal cells thus links *Paupar-Pax6* co-expression control and chromatin regulation to important neuro-developmental signalling pathways.

Our work also discovered a large chromatin domain downstream of the *Pax6* gene that displays clusters of increased long-range chromatin interactions with the *Paupar-Pax6* locus in E12.5 mouse neurons compared to ESCs. This region is further downstream from the previously defined *Pax6* distal DRR and the aniridia-associated breakpoints within the last intron of the downstream *ELP4* gene that are predicted to influence *Pax6* expression in affected individuals (Kleinjan, Seawright et al. 2001, Kleinjan, Seawright et al. 2006). However, it maps to an equivalent region in the human genome that contains multiple long-range enhancer-promoter looping interactions with the *Pax6* promoter, is characterised by an increase in enhancer-like chromatin modifications and is located with within the same self-interacting TAD as the *Paupar* and *Pax6* promoters in neurons (Bonev, Mendelson Cohen et al. 2017, Freire-Pritchett, Schoenfelder et al. 2017, Lu, Liu et al. 2020). Our results are thus consistent with a model in which developmentally regulated changes in distal chromatin architecture also play a role in CRE-promoter rewiring and the activation of *Paupar* and *Pax6* expression in the neuronal lineage.

*Paupar* lncRNA represses *Pax6* expression and can modulate *Pax6* splicing in a transcript- and cell type-dependent manner (Vance, Sansom et al. 2014, Singer, Arnes et al. 2019). Our results here report additional functions for the *Paupar* DNA locus in *Paupar-Pax6* expression control. NG Capture-C revealed reproducible *cis*-regulatory interactions between DNA sequences within the *Paupar* locus and the *Pax6* P0 promoter in neuronal cell types, consistent with phase III ENCODE data showing that *Paupar* overlaps five candidate CREs with an enhancer-like chromatin signature (Consortium, Moore et al. 2020). We found that TCF7L2 binds to a functional silencer element, termed CRE5, within the *Paupar* locus and that recruitment of the dCas9-KRAB chromatin repressor to the TCF7L2 motif led to an increase in both *Paupar* and *Pax6* expression. dCas9-KRAB recruitment is not predicted to directly block *Paupar* transcription (Gilbert, Horlbeck et al. 2014), as the targeted TCF7L2 motif lies approximately 2 kb downstream of the *Paupar* TSS, suggesting that the capacity of CRE5 to repress both *Paupar* and *Pax6* expression is independent of the *Paupar* transcript produced. Similarly, several studies have reported distinct roles for lncRNA transcripts and transcriptional regulatory elements within their DNA loci in gene expression control. The *Haunt* DNA locus contains several transcriptional enhancer elements that loop onto the *HoxA* gene to increase its expression whereas the *Haunt* transcript binds upstream of *HoxA* to induce a repressive chromatin state and block *HoxA* expression (Yin, Yan et al. 2015). *Tug1* DNA contains a CRE that represses multiple neighbouring downstream genes whilst the *Tug1* lncRNA acts in *trans* to regulate different genes (Lewandowski, Dumbovic et al. 2020).

The identification of shared *Paupar* and *Pax6* CREs also raises the possibility that the *Paupar* promoter may be able to control *Pax6* expression through CRE competition as described for several other lncRNAs. The promoters of the *Pvt1* lncRNA and neighbouring *Myc* oncogene compete for interactions with four shared enhancers. Silencing the *Pvt1* promoter using CRISPRi increased enhancer contacts with the Myc promoter and up-regulated Myc expression independent of the Pvt1 transcript (Cho, Xu et al. 2018). Similarly, the *Handsdown* locus interacts with several enhancers for the adjacent *Hand2* gene and regulates their usage during cardiac differentiation (Ritter, Ali et al. 2019). Our study provides significant new insights into the chromatin interactions, transcription factors and signalling pathways controlling *Paupar-Pax6* co-expression in the neuronal lineage and has general importance for understanding the wider role of lncRNA-mRNA transcription units in neuronal commitment, differentiation and function.

## Materials and Methods

### Plasmids

Individual sgRNAs were cloned into pX-dCas9-mod-KRAB to generate plasmids for CRISPRi as described in (Coe, Tan et al. 2019). Oligonucleotides used to clone sgRNAs targeting the TCF7L2 motif in CRE2, CRE3 and CRE5 are shown in S7 Table. The plasmid pCI-neo beta catenin S33Y was a gift from Bert Vogelstein (Addgene plasmid # 16519; http://n2t.net/addgene:16519; RRID: Addgene_l6519).

### Cell culture

Primary cortical neurons and cortical neural stem cells were prepared from CD1 mouse embryos (E14.5) in accordance with UK Home Office Guidelines as stated in the Animals (Scientific Procedures) Act 1986 using Schedule 1 procedures approved by the University of Bath Animal Welfare and Ethical Review Body. Cortices were dissected from embryonic brain and mechanically dissociated in PBS supplemented with 33 mM glucose, using a fire-polished glass Pasteur pipette.

#### Primary neurons

For preparation of differentiated cortical neurons (Cox, Choudhry et al. 2015), cells were plated into Nunc 90 mm petri dishes, previously coated with 20 μg/ml poly-D-lysine (Sigma), at a seeding density of 500 x10^5^ cells/ml. Neurons were cultured in Neurobasal medium (phenol red free) supplemented with 2 mM glutamine, 100 μg/ml penicillin, 60 μg/ml streptomycin and B27 (all from Gibco), and incubated at 37°C, in high humidity with 5% CO_2_. Under these growth conditions at 7 days *in vitro* (DIV) cells were non-dividing, had a well-developed neuritic network and were 99% β-tubulin III positive and <1% GFAP positive.

#### Primary neural stem cells

For preparation of cortical neural stem cells (Molina-Holgado, Rubio-Araiz et al. 2007), cells were plated into Nunc 90 mm petri dishes, previously coated with Cell Start (Gibco), at a seeding density of 500 x10^5^ cells/ml. Neural stem cells were cultured in StemPro NSC SFM composed of: Knockout D-MEM / F12; Glutamax (2mM); bFGF (20ng/ml); EGF (20ng/ml); StemPro Neural Supplement (2%); all from (Gibco). Under these growth conditions at 7 DIV cells were proliferative and were Nestin and Ki67 positive.

#### Cell Lines

N2A cells were grown in DMEM supplemented with 10% fetal bovine serum (FBS). For the mouse ESCs experiments E14Tg2A cells were maintained in GMEM supplemented with 10% FBS, 1xMEM nonessential amino acids, 2 mM glutamax, 1 mM sodium pyruvate, 100 mM 2-mercaptoethanol, and 100 units/ml LIF on gelatinised tissue culture flasks.

### Transfections and treatments

Approximately 3 x 10^5^ N2A cells were seeded per well in a 6-well plate for both plasmid DNA and esiRNA transfections. The following day, cells were transiently transfected using Lipofectamine 2000 (Invitrogen) following the manufacturer’s instructions. For knockdown experiments, cells were transfected with 1.5 μg MISSION® esiRNA (SIGMA-ALDRICH) targeting either *TCF7l2* (EMU010891) or Renilla Luciferase (EHURLUC) control and harvested 3 days later. 2μg pCI-neo beta catenin S33Y or empty vector were used in B-catenin S33Y overexpression experiments and cells were harvested 48 hrs post transfection. CRISPRi experiments were carried out as described in (Coe, Tan et al. 2019).

For Wnt and Bmp treatment, approximately 3 x 10^5^ N2A cells were seeded per well in a 6-well plate in either growth medium containing 50 ng/ μl Wnt3a (R&D Biosystems, 5036-wn) or in low serum medium (DMEM supplemented with 5% FBS) containing 0.1 ng/ μl BMP4 (Thermo Fisher, PHC9534) as described in (Szemes, Melegh et al. 2020). 0.02% BSA in PBS was used as a vehicle control. Cells were harvested for RNA extraction 72 hours later. Sequences of primers used for expression analysis are shown in Table S7.

### NG Capture-C

NG Capture-C libraries were prepared as described previously (Davies, Telenius et al. 2016). Briefly, approximately 2 x 10^7^ cells per sample were fixed with 2% formaldehyde for 10 min at RT, quenched by the addition of glycine and washed with PBS. Cell lysis was performed for 20 min on ice (10 mM Tris-HCl, pH8, 10 mM NaCl, 0.2% IGEPAL), lysed cells were then homogenized on ice and enzymatically digested overnight at 37°C with DpnII (New England Biolabs). The digested DNA was diluted and ligated with T4 DNA ligase overnight at 16°C. The following day, ligation reactions were de-crosslinked by Proteinase K (Thermo Scientific) addition and overnight incubation at 65°C. DNA extraction was performed by PCI and chloroform extraction followed by ethanol precipitation. DpnII digestion efficiency was confirmed by gel electrophoresis and quantified by real-time PCR – only 3C libraries with over 70% efficiency were used for the subsequent steps. Samples were sonicated to an average size of 200 bp using a Bioruptor Pico (Diagenode) and NEBNext Multiplex reagents and sequencing adapters were used to prepare sequencing libraries following the Illumina NEBNext DNA library prep kit instructions (New England Biolabs). Two rounds of capture using a pool of biotinylated oligos (IDT, see Table S7 for sequences) were performed on 1ug of each of the indexed libraries using the Nimblegen SeqCap EZ hybridization system. Library size was determined using the Tapestation D1000 kit and the DNA concentrations were measured on a Qubit 2.0 Fluorometer.

### Computation analysis of NG Capture-C data

Multiplexed NG Capture-C libraries were prepared from ESCs, NSCs, neurons and N2A cells (two biological replicates each) and 150 bp paired end sequencing was performed on the Illumina HiSeq 4000 (Novogene) to a total depth of approximately 500 million reads. The resulting fastq files from each of the eight replicates were combined using a Perl script. Raw reads were trimmed using trim_galore version 0.4.4 with parameter –paired. The trimmed paired-end reads were then combined using flash version 1.2.11 with the parameters --interleaved-output –max-overlap=200. The resulting fastq files of combined and uncombined reads were next merged into a single fastq file using the command cat. The fragments in the resulting fastq file were *in silico* digested into DpnII restriction enzyme digestion fragments using dpnII2E.pl. The resulting dpnIIE fragments were aligned to the mm10 genome using bowtie version 1.1.2 with parameters -p 1 -m 2 --best --strata --chunkmb 256 –sam. A set of DpnII fragments for the full mouse genome was produced from mm10.fa using gpngenome.pl. CCAnalyser3.pl was then run to compare a text file of the *Paupar, Pax6* P0, *Pax6* P1 and *Sox2* viewpoint coordinates with the *in silico* digested reads and genome to produce counts of the observed interactions with each viewpoint in each replicate. The output from CCAnalyser3.pl was analysed using the BioConductor r3Cseq package to determine statistical significance (p- and q-values) for the observed interactions between the viewpoints and each DpnII digestion fragment in each replicate. For each viewpoint, the resulting tables were combined into a table showing replicated significant fragments (significant in both replicates of at least one cell type) and these tables were used as the basis for subsequent analyses. Motif discovery was performed using the BiFa web tool at the Warwick Systems Biology Centre website (http://wsbc.warwick.ac.uk/wsbcToolsWebpage). 4Cin was used to generate three-dimensional models of *Paupar-Pax6* local chromatin architecture (Irastorza-Azcarate, Acemel et al. 2018).

### A statistical method for detecting meso-scale changes in chromatin conformation from NG Capture-C data (DeltaCaptureC)

We developed a new statistical method to detect changes in chromatin conformation based on significant clustering of neighbouring fragments from 3C-based data. This is available as a Bioconductor Software Package (10.18129/B9.bioc.deltaCaptureC). By binning NG Capture-C data and using permutation testing, this package can test whether there are statistically significant changes in the interaction counts between the data from two cell types or two treatments. To do this, read counts from the two biological replicates for each cell type were first combined. The counts for the four samples were normalised using DESeq2 function estimateSizeFactorsForMatrix() (Love, Huber et al. 2014) and the mean normalised count for both replicates in each cell type was determined. The difference between the two mean normalized counts was then calculated. This data was trimmed to a region of interest, 500kb up- and down-stream of the midpoint of viewpoint, binned to a fixed bin size of 1kb and then re-binned to 10kb (S2 Fig). This identified a large distal region of increased chromatin interactions in neurons (Fig 5 and S2 Fig). We observed that this a contiguous region of constant sign (negative) with a combined total absolute value of 308.8. The null hypothesis is that this sum arises by chance. We tested this hypothesis to detect statistical significance for continuous regions of constant sign in the following manner: we first excluded the region 50kb up- and downstream of the viewpoint and performed random permutation of the (non-viewpoint) 1kb bins. After each such permutation, data was re-binned to 10kb and each region was examined for constant sign. To do this, we computed its total absolute value and recorded the largest of these totals. If, after performing 1000 such random permutations, we observe fewer than 50 cases where the largest sum is 308.8 or greater, we have discovered a p-value for this region of less than 0.05. In this way, we can exploit co-localisation of differences with like sign to detect meso-scale chromatin remodelling from 3C-based data (Fig 5A and S2 Fig).

We then considered the region near the viewpoint. In this case it is important to note that raw NG Capture-C counts in this region strongly correlate with distance from the viewpoint and we are thus unable to perform arbitrary permutation to test for statistical significance. However, performing permutations which do not change this distance allowed us to test the null hypothesis that chromatin remodelling flanking the viewpoint was symmetric. To this end, we computed the difference between the sum in the region upstream of the viewpoint and downstream of it using the actual data. We then computed this difference after multiple symmetric permutations. Since there are 50 1kb bins in this region upstream and downstream of the viewpoint, there are 2^50^ permutations of this form giving us a sufficient number for permutation testing. In this way, we detected asymmetry in chromatin architecture in the neighbourhood of the *Paupar* and *Pax6* promoter viewpoints in each cell type (Fig 5A).

### RT-qPCR

RNA extraction was carried out using the GeneJET RNA Purification Kit (ThermoFisher) according to the manufacturer’s instructions with the addition of an on-column DNase digestion step using the RNase-free DNase Set (QIAGEN). Reverse transcription was performed using the QuantiTect Reverse Transcription Kit (Qiagen). 1 μg total RNA was used in each reaction. Quantitative PCR was carried out on a Step One Plus Real-Time PCR System using Fast SYBR Green Master Mix (Applied Biosystems).

### Chromatin Immunoprecipitation

ChIP experiments were performed as previously described using approximately 1 x 10^7^ N2A cells per assay (Vance, Sansom et al. 2014). Cross-linked chromatin was immunoprecipitated with either 5 μg anti-TCF4/TCF7L2 (Clone 6H5-3, #05-511, Millipore) or normal mouse control IgG (#12-371, Millipore) antibodies. qPCR primers used to amplify TCF7L2 motif containing sequences (CRE1-7) are shown in S7 Table.

## Data Availability

All multiplexed NG Capture-C DNA sequencing data for this project have been deposited in the NCBI GEO database (https://www.ncbi.nlm.nih.gov/geo/) under accession number GSE129697.

## Supporting Information Legends

### Figures

**S1 Fig.**
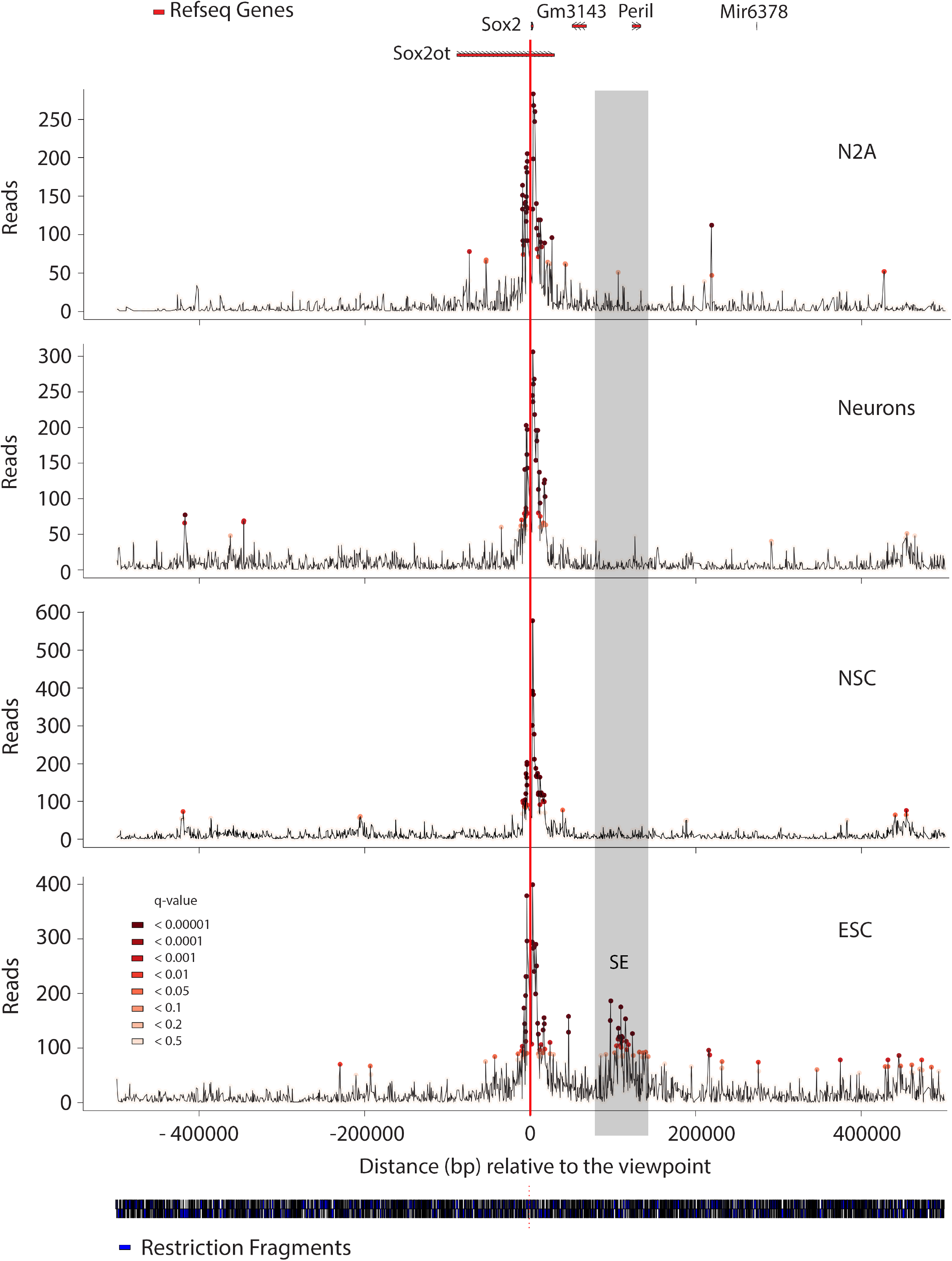
NG Capture-C identified ESC-specific chromatin looping interactions between the *Sox2* super-enhancer spanning the *Peril* locus and the *Sox2* promoter. NG Capture-C profiles displaying the *Sox2* promoter interaction count per DpnII restriction enzyme fragment in the indicated cell types. The red vertical line indicates the location of the *Sox2* promoter viewpoint. Significant interactions were determined using R3C-seq

**S2 Fig.**
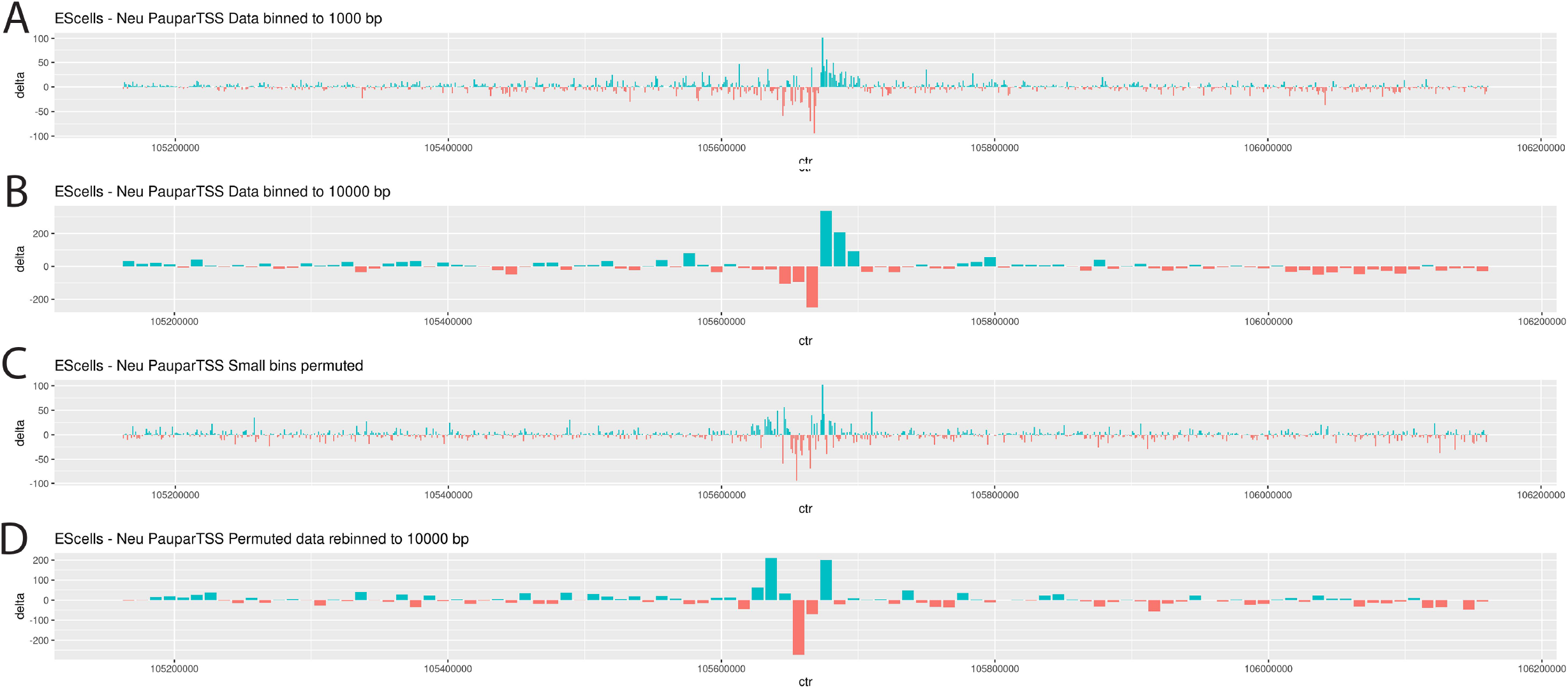
Increasing bin size facilitates the detection of chromatin changes between cell types. Differences in mean normalized NG Capture-C counts between neurons and ESCs for interactions with the *Paupar* viewpoint are plotted on the y-axis. X-axis shows position on chromosome 2 (GRCm38/mm10). Sequence data was permuted to assess specificity and statistical significance was calculated as described in Materials and Methods. (A) Difference in mean normalised interactions between ESCs and neurons binned to 1 kb. Negative values shown in red indicate increased interactions in neurons. (B) The same data binned to 10 kb. Note the emergence of the large red region approximately 350 kb downstream of *Pax6* and the asymmetric pattern near the viewpoint. (C) The 1 kb bins from (A) permuted. Bins further than 50 kb from the viewpoint are permuted at random. Bins closer than 50 kb are only permuted keeping their distance from the viewpoint. (D) The previous panel re-binned to 10kb. Notice that no large contiguous regions of constant sign appear, nor does the asymmetric pattern seen near the viewpoint in (B).

**S3 Fig.**
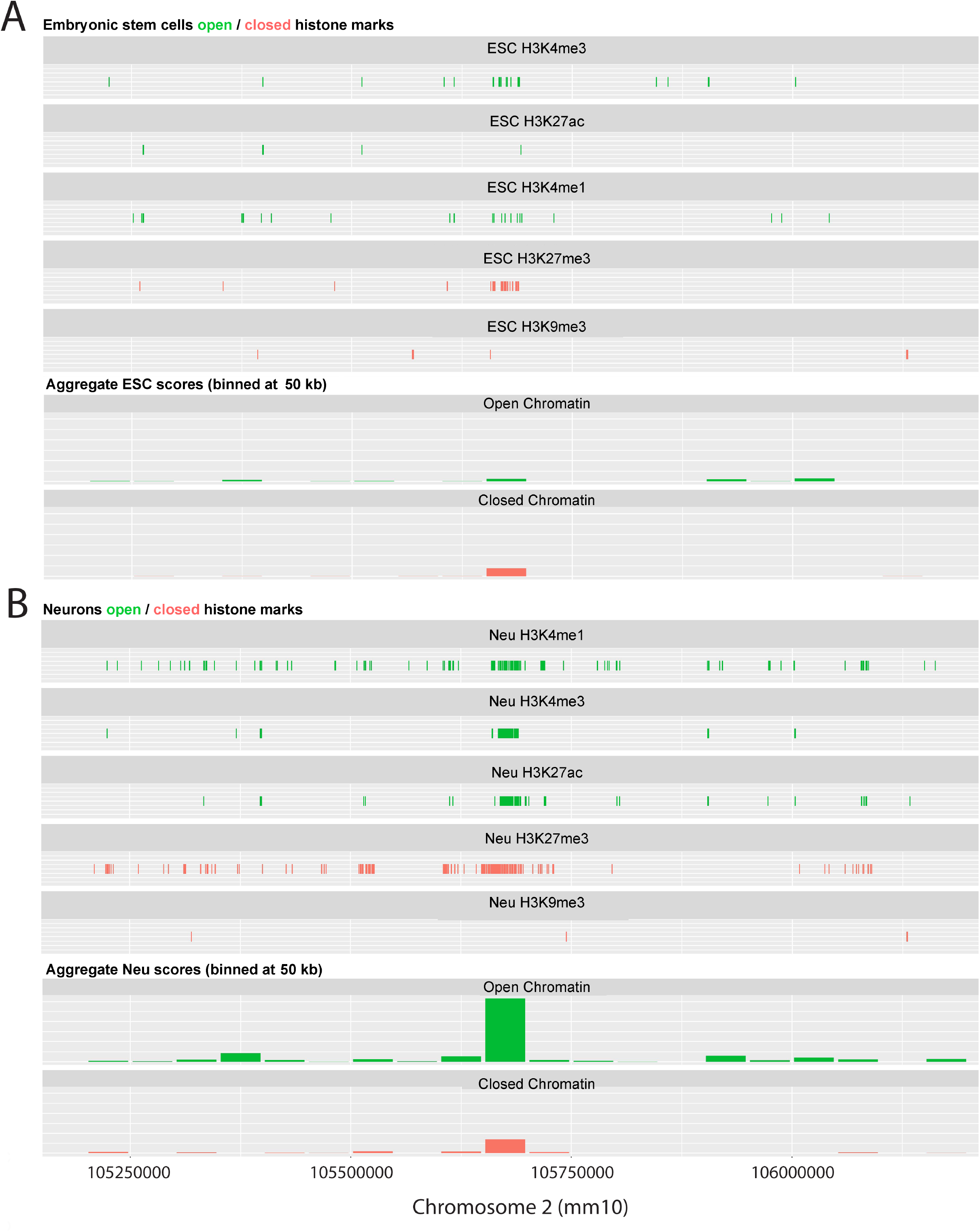
An increased frequency of chromatin interactions with the *Paupar* and *Pax6* promoters in neurons correlates with elevated levels of open compared to closed histone modifications. ENCODE Project ChIP-seq data mapping the location of open (H3K4me1, H3K4me3 and H3K27ac) and closed (H3K27me3 and H3K9me3) ChIP-seq peaks in ESCs (A) and E12.5 mouse forebrain tissue (B) across approximately 1MB genomic sequence surrounding the *Paupar-Pax6* locus (He, Hariharan et al. 2020). Individual peaks of less than 1000 bp are shown at 1000 bp long for visibility reasons. Aggregated data represent summed and binned scores from the individual tracks.

## Tables

**S1 Table. Determination of ligation and capture frequencies for NG Capture-C experiment**

**S2 Table. Number of unique interactions with each promoter viewpoint**

**S3 Table. Genome coordinates of significant replicated fragments for each viewpoint (Mouse GRCm38/mm10)**

**S4 Table. Interactions between the indicated viewpoints and reporter fragments that overlap the *Paupar* genomic locus**

**S5 Table. Number of replicated interactions in different cell types**

**S6 Table. BiFa analysis identification of TCF7L2 motifs**

**S7 Table. Oligonucleotides**

## Acknowledgements

This project has been funded by a Biotechnology and Biological Sciences Research Council grant to KWV (BB/N005856/1; KWV, IP, MS). We thank Dr Karim Malik (University of Bristol) for providing recombinant Wnt3a and Bmp4.

